# PeSA: A Software Tool for Peptide Specificity Analysis

**DOI:** 10.1101/760140

**Authors:** Emine Topcu, Kyle K. Biggar

**Affiliations:** Institute of Biochemistry and Department of Biology, Carleton University, 1125 Colonel By Drive, Ottawa Ontario, K1N 5B6 Canada

**Keywords:** Peptide specificity, Motif, Oriented Peptide Array Library, Permutation Array, Position-Specific Scoring Matrix

## Abstract

The discovery of molecular interactions is crucial towards a better understanding of complex biological functions. Particularly protein-protein interactions (i.e., PPIs), which are responsible for a variety of cellular functions from epigenetic modifications to enzyme-substrate specificity, have been studied extensively over the past decades. Position-specific scoring matrices (PSSM) in particular are used extensively to help determine interaction specificity or candidate interaction motifs. However, not all studies successfully report their results as a candidate interaction motif. In many cases, this is the result of a lack of analysis tools for simple analysis and motif generation. Peptide Specificity Analyst (PeSA) is developed with the goal of filling this gap and providing an analysis software to aid peptide array analysis and subsequent motif generation. PeSA utilizes two models of motif creation: (1) frequency-based using a peptide list, and (2) weight-based using a quantified matrix. The ability to generate motifs effortlessly will make analyzing, interpreting and sharing peptide specificity study results in a simple and straightforward process.

**GRAPHICAL ABSTRACT:** 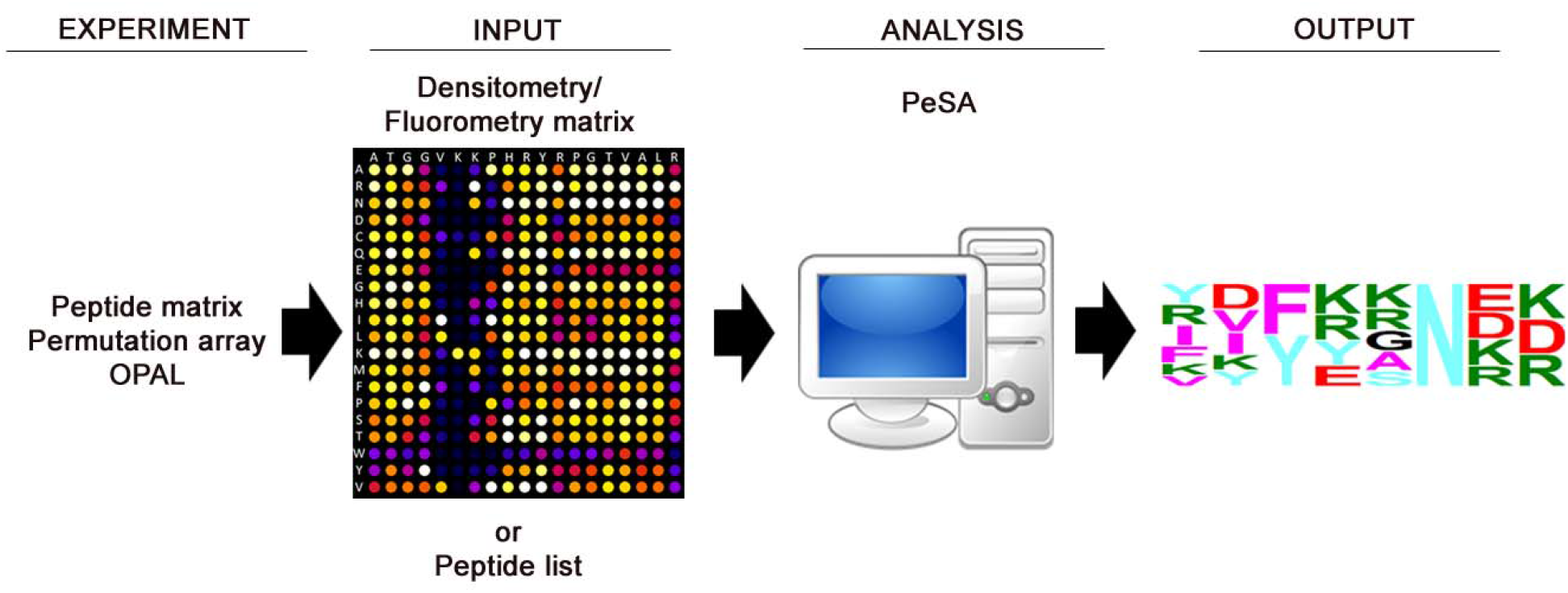

**HIGHLIGHTS:**

- Biological motifs are widely used representations for peptide specificity analysis.
- PeSA populates a list of peptides matching a set threshold from a quantified matrix.
- Frequency-based motif using a peptide list to spot residue patterns.
- Use of quantified matrices to create weight-based motifs using residue positions.

## 1. INTRODUCTION

The discovery of molecular interactions is crucial towards a better understanding of complex biological functions. Particularly protein-protein interactions (i.e., PPIs), which are responsible for a variety of cellular functions from epigenetic modifications to enzyme-substrate specificity, have been studied extensively over the past decades (Arenkov et al., 2000; Arifuzzaman et al., 2006; Arrowsmith et al., 2012; Lassle and Blatch, 1999; Rual et al., 2005). Since not all molecular interactions, especially transient or low-stoichiometry interaction events, can be witnessed in vivo being able to identify or predict a possible interaction is of essential importance.

One modern approach used to identify and characterize PPI events relies on testing binding sites based on sequence similarity to known binding sites, creating a PPI motif that can be used to predict novel PPI events(Lanouette et al., 2015; Schuhmacher et al., 2015). Various methods have been developed throughout the years to predict and detect PPIs. In silico methods (e.g., PIPE (Pitre et al., 2006), STRING (Franceschini et al., 2013), FpClass (Kotlyar et al., 2014), and PRISM (Baspinar et al., 2014)) can generate high number of predictions using different characteristics of proteins, such as structure, domains, or similarities to other proteins with known interactions. Validation of these predictions can be done via medium and high-throughput assays such as protein-fragment complementation assays (Li et al., 2017, 2019; Primeau et al., 2002), yeast two-hybrid screens (Ito et al., 2001; Uetz et al., 2000; Weimann et al., 2013), nucleic acid programmable protein arrays (Rolland et al., 2014; Yu et al., 2018) and peptide arrays (Levy et al., 2011; Rathert et al., 2008). In this way, peptide arrays have been used as a cost effective and medium-throughput technique used to run parallel experiments on vast number of peptides (Lanouette et al., 2015; Rathert et al., 2008). The use of systematically mutated (i.e., permutated) peptide arrays in particular has been used frequently in PPI and enzyme-substrate specificity analysis (Dhayalan et al., 2011; Kudithipudi et al., 2014; Lanouette et al., 2015; Schuhmacher et al., 2015; Weirich et al., 2016). These permutation arrays are based on the systematic alternation of a single residue to all other amino acids at each position throughout a peptide sequence, resulting in a 2-dimensional matrix representing the substrate-space tolerable for an interaction to occur in vitro. By studying the impact of single amino acid change in a peptide sequence, the individual role of each amino acid can be used in making a prediction on the PPI or enzyme-substrate specificity(Lanouette et al., 2015).

Position-specific scoring matrices (PSSM) in particular are used extensively in peptide specificity studies (Creixell et al., 2015; Liu et al., 2013; Murakami et al., 2017; Zhu et al., 2005). However, not all studies successfully report their results as a candidate interaction motif. In many cases, this can be the result of a lack of a suitable tool for simple analysis and motif generation. The development of a motif manually or using tools that are not designed specifically for motif generation can be cumbersome and time consuming. Utilizing different cut off thresholds for decision making and running comparisons can be difficult to achieve, as well as error prone due to repetition of manual calculations and data manipulations. <u>Pe</u>ptide <u>S</u>pecificity <u>A</u>nalyst (PeSA) is developed with the goal of providing analysis software to aid peptide array analysis and motif generation.

PeSA has been designed to facilitate the interpretation of peptide array results. Developed using the enzyme-substrate identification literature for lysine methylation as a prototypical example of an experimental workflow, PeSA is also applicable to other protein-protein interactions and enzyme specificity analysis. Additionally, it has been expanded to also support results of degenerate Oriented Peptide Array Library (OPAL) studies (Wei et al., 2018). Using the number-of-occurrences, or the values of quantified data measured by the study, PeSA derives an amino acid motif from a peptide list or peptide array as a means of PSSM or position weight matrix (PWM). With a goal of developing an easy to use user-interface for occasional or non-expert users, PeSA provides an environment to view and visually compare results of an experiment with an adjustable cut-off threshold available for dynamic motif generation. PeSA can also be used to annotate a motif not only for the output of the study, but also for the input of the study to detect any existing bias and interpreting the results accordingly.

Current version of PeSA (v.1.5) provides a working environment to analyze data and create an amino acid sequence motif using different types of inputs, such as a list of peptides or quantified peptide matrix. The approach PeSA is taking is to minimize the manual editing or manipulation of the data by the user and to use the files that are generated for or by other software tools commonly used in the industry, such as Excel file output of ImageJ software or text files that are fed into peptide synthesizers. PeSA also allows the user to save the data as a PeSA project file or export as an Excel file for further manipulation if required.

Our prototypical framework of which the utility of PeSA is demonstrated is the use of peptide arrays to study lysine methylation, however the implementation of PeSA expands broadly beyond this example. Post translational modification (PTM) is one of the reversible and dynamic mechanisms that functions to increase the functional diversity of proteins beyond their genetic code. As such, PTMs directly contribute to influence a wide range of cellular processes through the regulation of protein function, stability and location. Lysine methylation is a commonly studied PTM (Kudithipudi et al., 2014, 2012; Liu et al., 2013; Rathert et al., 2008; Weirich et al., 2016). Both histone and non-histone proteins can go through lysine methylation (Dhayalan et al., 2011; Rayasam et al., 2003). Lysine methylation of histone proteins result in changes in genetic expression, whereas methylation of non-histone proteins can cause changes in cellular activity: cell growth, DNA repair, protein synthesis (Biggar and Li, 2014). Understanding the mechanism and making reasonable predictions on which lysine residues will go methylation is important in understanding biological functions and can be helpful in drug development. The example usage of PeSA for various lysine methylation studies discussed in this study are an indicator of its added value to current studies in such an important area of modern research. PeSA was developed as an opensource, easy to use tool designed to help researchers to analyze peptide sequences. The ability to generate motifs effortlessly will make analyzing, interpreting and sharing peptide specificity study results a simple and straightforward process.

## 2. MATERIAL AND METHODS

### 2.1 Design and Implementation

PeSA utilizes two models of motif creation: (1) frequency-based, and (2) weight-based. In the former frequency-based model, weight of each amino acid residue in PSSM is calculated by number of occurrences of an amino acid at a specific position divided by total number of peptides. If the frequency calculated is higher than the selected threshold, it is displayed in the motif. The height of the letter representing an amino acid is proportional to the frequency calculated. The amino acids of which the frequencies fall below the user-defined threshold are displayed as a single character ‘X’. In contrast, for the latter weight-based model, the quantification values computed by the study results are used to determine the height of the letter representing an amino acid in the motif, rather than the mere frequency.

#### 2.1.1 User-supplied peptide list

As its simplest from, PeSA accepts a user-supplied peptide list to create a conserved sequence motif. The frequencies that will be used to create a frequency-based motif as described above.

#### 2.1.2 Peptide arrays

PeSA is also able to accept sequences from a peptide array matrix and associated raw quantified matrix (i.e., densitometry, fluorometry, etc.) to create the conserved sequence motif. The quantification matrix is automatically normalized by identifying and using the ratio of the maximum numerical value within the matrix, or a manually defined value by the user. Peptides that have a normalized value greater than a user-defined threshold ratio are then selected to be used in the creation of a frequency-based motif.

#### 2.1.3 Permutation arrays

Permutation arrays have been widely used to systematically explore position-specific tolerable residue (i.e., amino acid) substitutions in protein-interaction and enzyme-substrate experiments. For these experiments, a weight-based model is adopted for the generation of sequence motifs. The user-supplied experimental results of the quantification matrix determine the weight of the modified residues at each position after normalization. Normalization of the experimental results occurs in a manner similar to that described above, using the numerical value of the wildtype sequence within the row or column. When the permutation array is formatted with each vertical column representing a residue mutation at a specific position, PeSA will automatically normalize the data in a column-wise manner. For column-wise normalization, each value within each column of the quantified matrix is divided by the value of the wildtype sequence (i.e., normalization value) within that column. PeSA will automatically identify the axis holding the permutation amino acid values, the wildtype sequence in the opposite axis, and the appropriate quantified value within each column/row. Given that there is no repetition of amino acids in the permutation axis, this information is used to make the axis determination. In the event that the wildtype sequence also does not have any residue repetition, preventing PeSA from recognizing the proper axis designations, the user has the ability to manually identify which axis represents the wildtype sequence information. The normalization value any specific column/row is determined by the peptide spot within the array equivalent to the wildtype sequence (i.e., the residue permutation which is the same as the wildtype sequence). If PeSA is unable to identify an appropriate normalization value for a specific column, then the average of other columns’ normalization values is used. Transposed row-based residue mutations are handled in a similar manner.

#### 2.1.4 Oriented peptide array libraries

Lastly, when highly degenerate OPAL arrays are used for analysis, a weight-based model is adopted for the generation of sequence motifs similar to that described for permutation array analysis above. As there is no wildtype sequence for normalization for an OPAL array, the maximum value at each row or column is automatically identified and is used as the normalization value to determine the final weight of each residue at a specific position.

### 2.2 PeSA software details

PeSA has been implemented in C# language targeting .NET Framework platform (v.4.5+) using Microsoft Visual Studio Community 2017 (Microsoft Corporation, v.15.9.11). The software has two main components: (1) PeSA.Engine is the core component completing the analysis and motif creation. Having a separate component layer provides reuse of the business logic by other platforms with only user interface implementation. (2) PeSA.Windows is the user interface application built using the Windows Forms Library.

PeSA uses external packages from NuGet package repository (https://www.nuget.org): (1) Newtonsoft.Json (Newton-King, 2019) is used for serialization and deserialization of models in JSON format. (2) EPPlus (Källman, 2019) is used for importing or exporting Excel files. The source code for the project can be downloaded from www.github.com/EmineTopcu/PeSA repository. The code can be compiled by Visual Studio 2015 (or later versions) IDE (Integrated Development Environment) of Microsoft.

### 2.3 PeSA features

Current version of PeSA (v.1.5) can create an amino acid sequence motif using; (1) a peptide list, (2) quantified peptide array, (3) permutation array or (4) OPAL, with a dedicated user-interface for each type of analysis. Implementation and use of each analysis are described below.

#### 2.3.1 Create sequence motif from peptide list

This module allows the user to create a motif based on frequency of amino acids at each position. Each amino acid is represented as its single letter acronym, relative to its frequency within the provided peptide list. The color of each amino acid can be modified through Settings → Motif Settings. The screenshot of the user interface and the result motifs are displayed in Figure 1.

**Figure 1.**
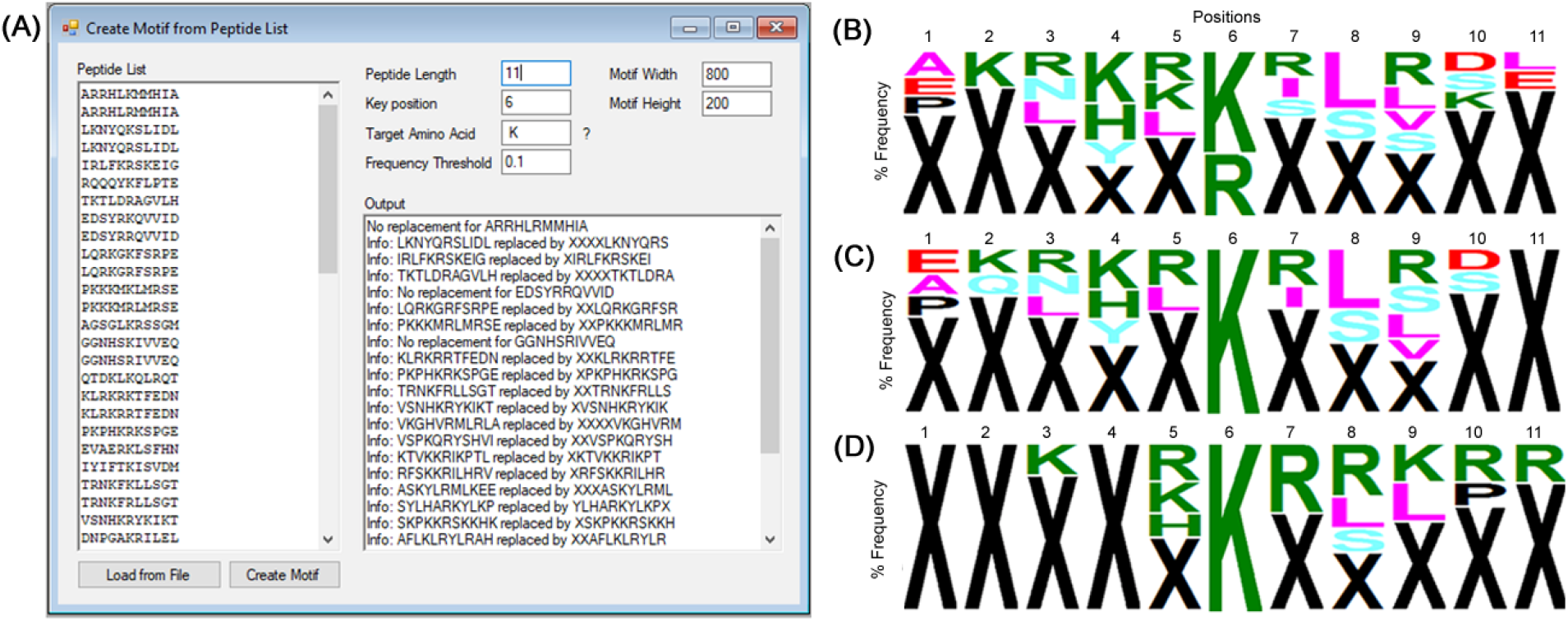
Creating a sequence motif from peptide list user interface. (A) The user interface with a sample input peptide list is shown. (B) Single motif generated without entering Target Amino Acid. The colors representing letters of amino acid residues can be set within the Settings of PeSA. Current settings use chemical properties of the residues. Acidic polar (DE): Red; Basic polar (RHK): Green; Neutral polar (NQSTY): Blue; Nonpolar (AILMFWV): Pink; Special (CGP): Black (C) Main motif generated if the Target Amino Acid information is entered. (D) The second motif generated by shifting the peptides so that target amino acid residue resides at the key position. The second motif is generated only if the Target Amino Acid is entered. The shifted peptides used to generate this motif is displayed on the Output window of (A).

##### Input

Motif generation from a peptide list feature requires a list of peptides, each peptide on a separate line. The list can be entered manually or loaded directly from a file. Accepted file types are text files (*.txt), Excel files (*.xls, *.xlsm, *.xlsx) and comma delimited files (*.csv).

##### Peptide length

This is an optional parameter. If left blank, the length of the first peptide in the list will be accepted as the peptide length. Entering the peptide length allows users to reuse a file fed to a peptide generation machine. In cases where there are extra characters appended to each peptide as required by the peptide generation machine, these additional characters can be ignored by PeSA by entering the “peptide length” parameter.

##### Key position

If not entered, key position is calculated as the middle position of a peptide. This parameter is used only if Target Amino Acid parameter is entered.

##### Target amino acid

The difference between the motifs created with and without a target amino acid can be viewed in Figure 1B and 1C/D, respectively. If no target amino acid is entered, only one motif is created and every position in the peptide is handled the same way. In the presence of a key position and target amino acid, a motif is created by using frequencies of the amino acids only in the peptides with that specific amino acid in the key position. Also, a secondary motif is generated for the rest of the peptides, if any of the peptides not included in the first motif can be shifted left or right to bring the target amino acid to the key position.

##### Frequency threshold

This parameter is the minimum percentage to be displayed in the motif. For example, with a threshold of 0.1, an amino acid is represented at a specific position in the motif only if it exists in at least 10% of the peptides in the list at the same position.

##### Motif width and height

Dimensions of the motif to be generated can be entered here as number of pixels. The default values can also be updated through Settings → Motif Settings.

##### Output

The output box displays warnings or error messages, e.g. nonstandard amino acid abbreviations, invalid characters in the peptide list, or inconsistent peptide lengths. The details about how the second motif is generated by sequence shifting are also displayed in this window. Motifs created are displayed in a separate window, which can be saved as an image file or copied to the clipboard.

#### 2.3.2 Peptide array analysis

Peptide Array Analysis module allows the user to determine the list of peptides from a peptide array to feed to the motif generator discussed in section 2.3.1. Based on the quantification values imported for the peptide array, the normalize by and threshold values set, the peptides that are accepted as modified are marked with gray background for easy identification. The user has the capability of saving all the imported, modified and generated data as a PeSA project file to open later or exporting them to an Excel file for further manipulation/analysis outside of PeSA. The screenshot of the user interface and the result motifs are displayed in Figure 2.

**Figure 2.**
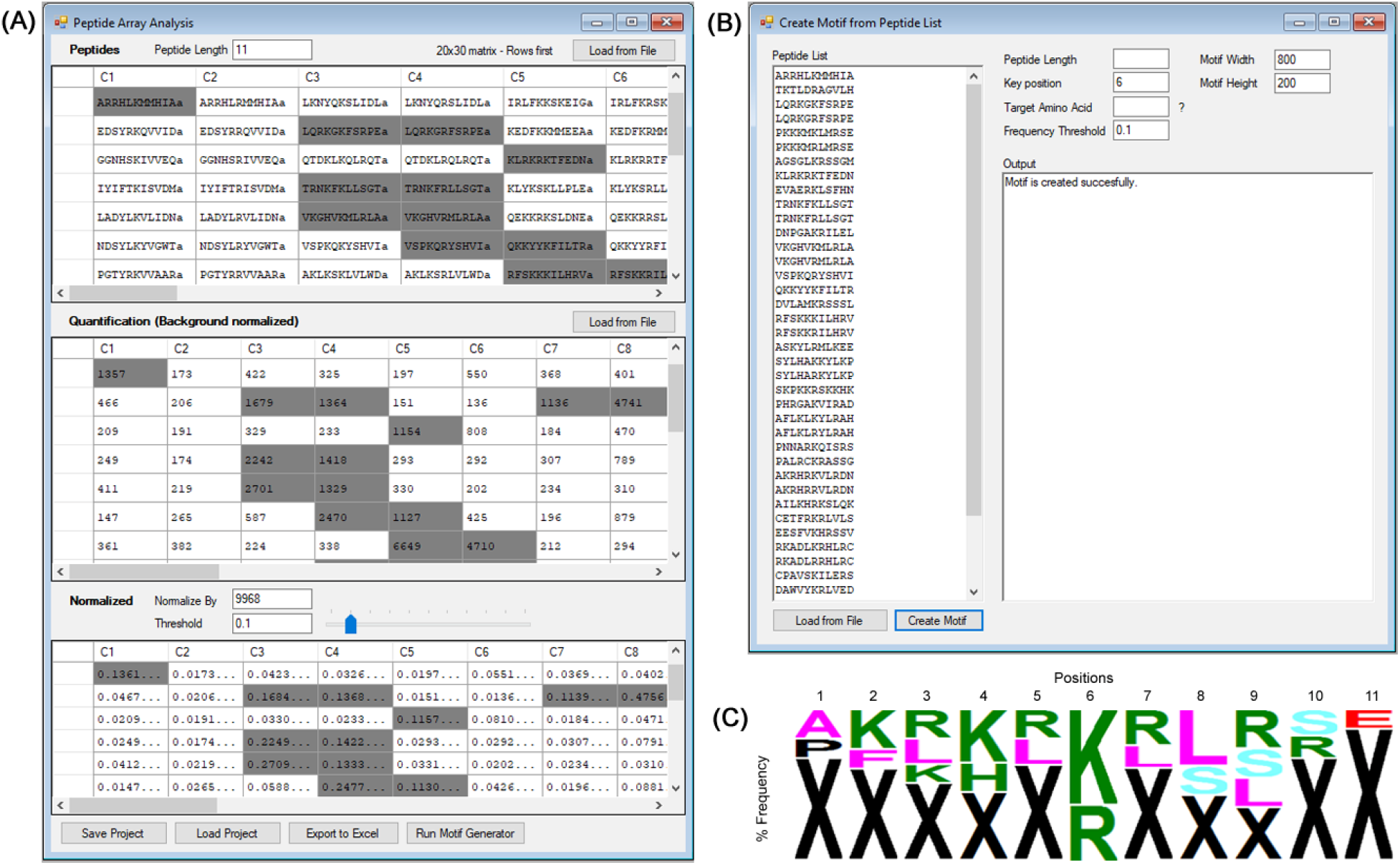
Peptide Array Analysis user interface. (A) The user interface with a sample input peptide matrix and corresponding quantification matrix is shown. The normalized matrix is generated real time, using the ‘normalize by’ and ‘threshold’ values that can be modified by the user. (B) Running motif generator through the Peptide Array Analysis user interface will run the motif generator user interface with the peptide list that meets the criteria. (C) Frequency-based motif generated for the corresponding peptide list. The colors of the amino acid residues are set as: Acidic polar (DE): Red; Basic polar (RHK): Green; Neutral polar (NQSTY): Blue; Nonpolar (AILMFWV): Pink; Special (CGP): Black.

##### Peptide sequence (input)

The peptide matrix can be pasted as clipboard data from Excel or loaded from a file. Accepted file formats are Excel files (*.xls, *.xlsm, *.xlsx) and comma delimited files (*.csv). The dimensions of the matrix are determined automatically based on the loaded/copied matrix. A list of peptides can also be imported in text format, with each peptide on a different line. In that case, the matrix is generated by PeSA. The dimensions of the matrix and whether the generation is on columns first basis or rows first basis can be set through Settings → Peptide Array Settings. The peptide grid also provides search functionality for partial or full peptide sequence within the matrix.

##### Quantification value (input)

The quantification matrix generated by other software needs to be imported to PeSA. It can be directly copied from Excel clipboard or loaded from a file. Accepted file formats are Excel files (*.xls, *.xlsm, *.xlsx) and comma delimited files (*.csv).

##### Peptide length (parameter)

This is an optional parameter, identical to the peptide length in section 2.3.1.

##### Matrix dimensions (display only)

The dimensions of the matrices are determined automatically if the peptides are copied or uploaded in matrix format. If a peptide list is loaded instead, the dimensions of the matrix need to be entered through Settings → Peptide Array Settings.

##### Normalize By (parameter)

By default, the maximum value in the quantification matrix is taken as the normalize by value. To create the normalized matrix, every value in the quantification matrix is divided by the normalize by value.

##### Threshold (parameter)

The threshold value is used to determine whether a peptide in the peptide matrix is accepted as modified or not. Any value in the normalized matrix that is equal to or more than the threshold value is accepted as modified, and the corresponding peptide is added to the peptide list which is to be fed to the motif generator. The threshold can be set to any value between 0 and 1 using the slider, or by entering any numeric value using the entry box.

#### 2.3.3 Permutation array analysis

Permutation Array Analysis module allows the user to create a weight-based motif using the residues at a specific position that provide a quantification value greater than a provided threshold. The peptides that are accepted as modified are displayed with gray background as in the other modules of the software for easy identification. The input and output data can be saved as a PeSA project file for later access or exported as an Excel file for further manipulation. The screenshot of the user interface and the result motifs are displayed in Figure 3A-3B.

**Figure 3.**
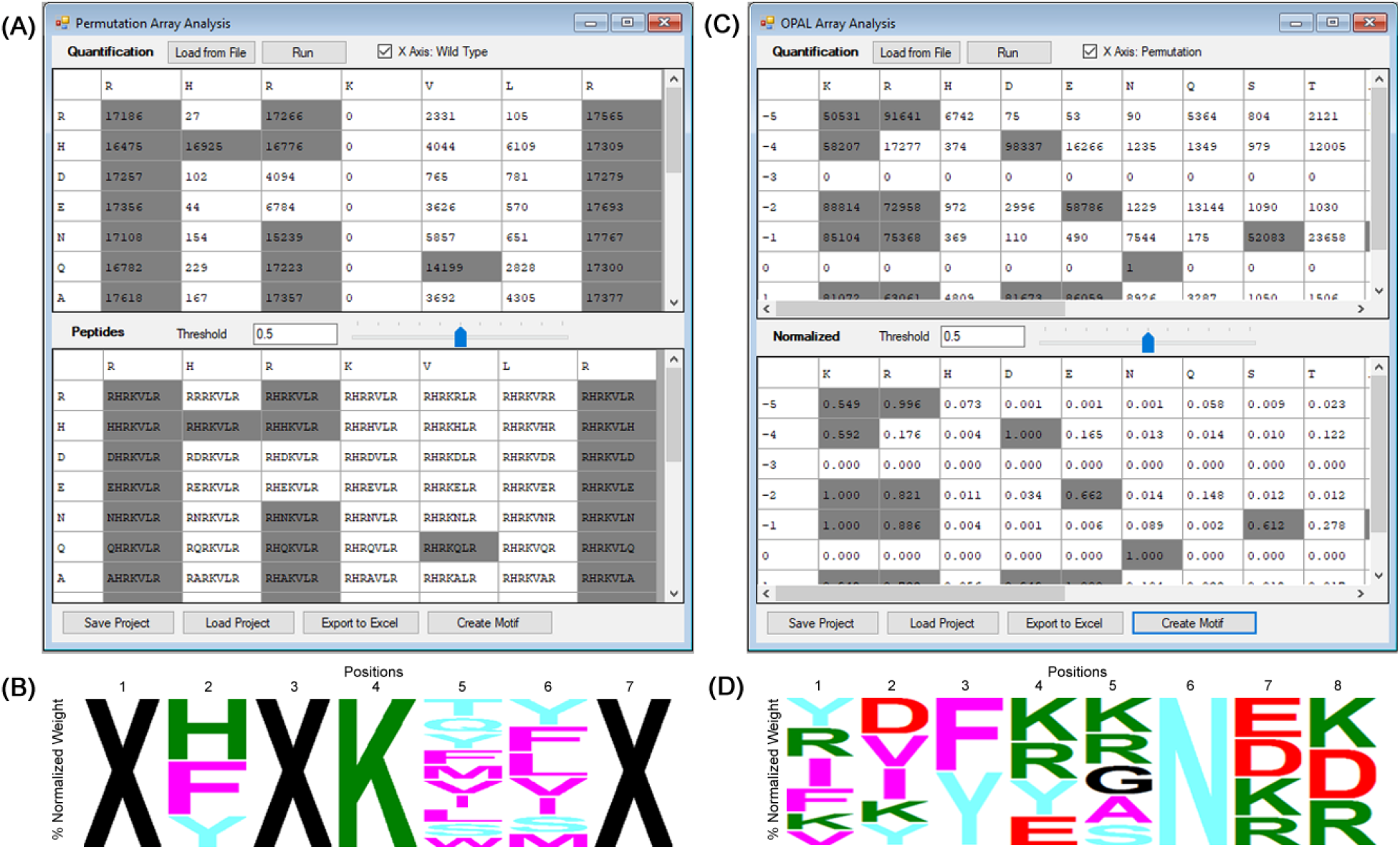
Permutation Array and OPAL Analysis user interface. (A) The user interface with a sample permutation array and corresponding quantification matrix is shown. The normalization takes place by the quantification values divided by the wildtype value within the corresponding row or column. Peptides with normalized values greater than the user set threshold are displayed with gray background. (B) Weight-based motif generated for the corresponding peptide list. The colors of the amino acid residues are set as: Acidic polar (DE): Red; Basic polar (RHK): Green; Neutral polar (NQSTY): Blue; Nonpolar (AILMFWV): Pink; Special (CGP): Black. (C)The user interface with a sample quantification of an OPAL array and corresponding normalized matrix is shown. The normalization takes place by the quantification values divided by the maximum value within the corresponding row or column. Peptides with normalized values greater than the user set threshold are displayed with gray background. (D) Weight-based motif generated for the corresponding peptide list.

##### Quantification value (input)

The quantification matrix containing the wild type sequence on one axis (i.e. as row or column header), the permutation values on the other axis, and background normalized quantification values as the rest of the matrix needs to be imported to PeSA. It can be directly copied from Excel clipboard or loaded from a file. Accepted file formats are Excel files (*.xls, *.xlsm, *.xlsx) and comma delimited files (*.csv).

##### X-Axis: wild type (input)

PeSA has the ability to determine which axis contains the wild type sequence. In case the wild type sequence does not provide a cue, the user can enforce the appropriate axis selection.

##### Peptide matrix (output)

The peptides generated by using the wild type sequence and the permutation values are displayed in the matrix format, by which the user can determine the accuracy of matrix orientation. The background color of the cells is set to gray if the normalized values are greater than the set threshold. The normalized values (not displayed) are calculated by dividing the quantification values by the wild type sequence value of the corresponding row or column.

##### Threshold (parameter)

The threshold value is used to determine whether a peptide in the permutation matrix is accepted as modified or not. Any peptide with a normalized value that is equal to or more than the threshold value is accepted as modified. The threshold can be set to any value between 0 and 1 using the slider, or by entering any numeric value using the entry box.

#### 2.3.4 OPAL array analysis

OPAL Array Analysis module is an extension of PeSA to manage oriented peptide array libraries where there is no wild type sequence, but each location of the matrix differs by one residue concentration at a specific position being higher than the rest. A weight-based motif is generated using the normalized values of the quantification value greater than the provided threshold. The peptides that are accepted as modified are displayed with gray background as in the other modules of the software for easy identification. As with the other modules, the input and output data can be saved or exported as an Excel file. The screenshot of the user interface and the result motifs are displayed in Figure 3C-3D.

##### Quantification value (input)

The quantification matrix containing the permutation values on one axis (i.e. as row or column header), and background normalized quantification values as the rest of the matrix needs to be imported to PeSA. The values on the column or row header opposite to permutation values are ignored. The quantification matrix can be directly copied from Excel clipboard or loaded from a file. Accepted file formats are Excel files (*.xls, *.xlsm, *.xlsx) and comma delimited files (*.csv).

##### X-Axis: permutation (input)

PeSA has the ability to determine which axis contains the permutation values. In case the header values do not provide a cue, the user can enforce the appropriate axis selection.

##### Normalized matrix (output)

The normalized values are calculated by dividing the quantification values by the maximum value in the corresponding row or column. The background color of the cells is set to gray if the normalized values are greater than the set threshold.

##### Threshold (parameter)

The threshold value is used to determine whether a peptide in the OPAL matrix is accepted as modified or not. Any peptide with a normalized value that is equal to or more than the threshold value is accepted as modified. The threshold can be set to any value between 0 and 1 using the slider, or by entering any numeric value using the entry box.

## 3. RESULTS AND DISCUSSION

PeSA uses quantified array results as raw input for further sequence analysis.The examples displayed in this article use the Protein Array Analyzer plugin (Carpentier, 2014) implemented using ImageJ software (Rasband, 2015) to convert autoradiography images to quantified raw densitometry values. To calculate the number of matches using PeSA, raw data used to generate the motif was utilized. For the sake of replicability, raw data of the motif is included in the Excel export files as well as the motif image (Supplemental tables S1-S10).

### 3.1 NSD1 methyltransferase substrate specificity analysis

To explore the substrate specificity of the NSD1 lysine methyltransferase, one notable study utilized a permutation array with the wildtype sequence H3(31-49) with K36 as the target lysine was methylated (Kudithipudi et al., 2014). The quantification matrix created by the densitometry of autoradiographic image of the same experiment was fed into PeSA (Figure 4), 2.3.3 Permutation array analysis feature, and the motif in Figure 4C was generated. In the same study, Kudithipudi and collegues (2014) reported that two non-histone proteins ATRX (K1033) and U3 (K189) were weakly methylated by NSD1, while p65 (K218 and K221) did not show any methylation. PeSA-generated motifs were compared to amino acid sequences of these candidate methylation sites, between −5 and +7 positions (Figure 4D). ATRX protein displayed a 100% match to threshold value of 0.60. U3 protein displayed a 100% match to a threshold value of 0.44, dropping to only 92.31% (corresponding to a single amino acid mismatch) at a 0.50 threshold. The p65 protein, targeting both K218 and K221 positions were found to have a 69.23% residue match at 0.50 threshold. Accepting a 0.5 threshold for analysis and a cut-off point of maximum one mismatch for prediction, PeSA would predict ATRX (K1033) and U3 (K189) to be methylated, and no methylation of p65 protein. These predictions agree with the predictions of Kudithipudi et al., as well as their experimental validation experiments.

**Figure 4.**
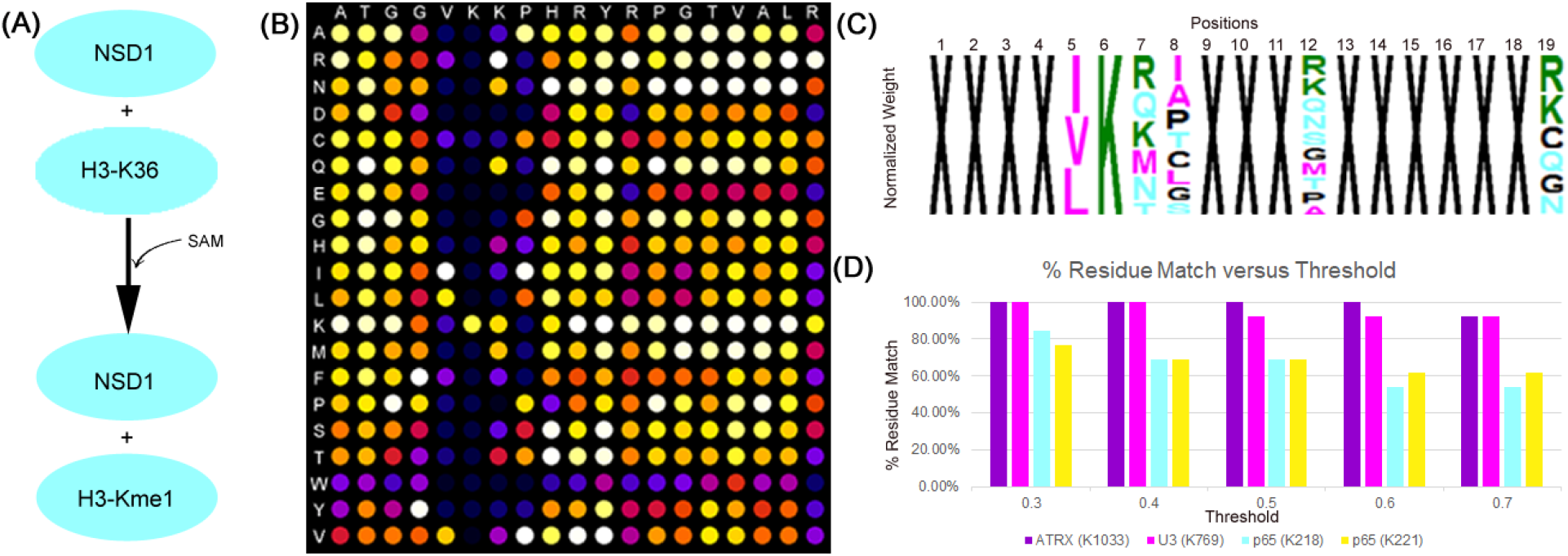
Analysis of NSD1 substrate analysis study (Kudithipudi et al., 2014) with PeSA. (A) NSD1 causes methylation of H3 protein at K36 position. (B) Densitometry analysis of a permutation array using H3 (31-49) sequence. (C) Specificity matrix created by PeSA using the quantification matrix created using the same permutation array results using a threshold value of 0.5. The colors of the amino acid residues are set as: Acidic polar (DE): Red; Basic polar (RHK): Green; Neutral polar (NQSTY): Blue; Nonpolar (AILMFWV): Pink; Special (CGP): Black (D) Scoring of the ATRX (K1033), U3 (K769)m and p65 (K218, K221) proteins using the % residue match at different thresholds.

### 3.2 SMYD2 methyltransferase substrate specificity analysis

A similar comparison was done for the study results of SMYD2 substrate specificity (Lanouette et al., 2015). Using their SPOT peptide array analysis of p53 (K370), a PeSA motif was generated (Figure 5A). The PeSA-generated motif was then compared to known substrates (p53 (K370), Erα (K266), Rb (K810), Rb (K860), and HSP90 (K615)), as well as the substrates predicted by Lanouette et al. (SIX1 (K51), SIN3B (K354), and DHX15 (K515)) (Figure 5B and 5C). The PeSA motif generated displayed notable amino acid specificity for positions −1 to +3. The PeSA match was calculated for each of the peptide sequences in question using different thresholds, for full motif (Figure 5B) as well as partial motif covering the [-1, +3] positions (Figure 5C).

**Figure 5.**
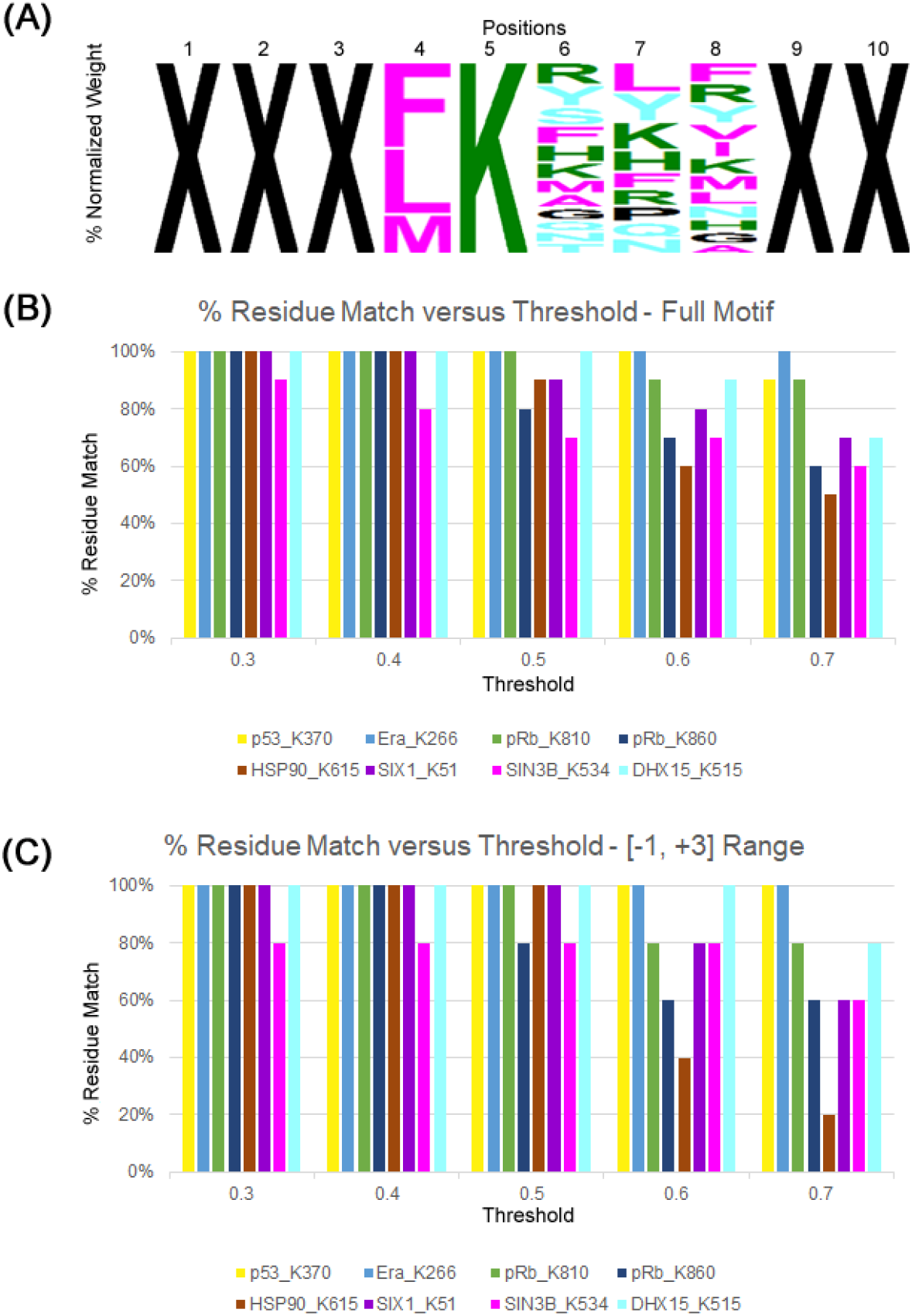
Analysis of SMYD2 substrate specificity study (Lanouette et al., 2015) with PeSA. (A) With a threshold value of 0.5, highest specificity for SMYD2 can be seen between −1 and +3 positions based on the permutation array created for peptide sequence of SSHLKSKKGQ. The colors of the amino acid residues are set as: Acidic polar (DE): Red; Basic polar (RHK): Green; Neutral polar (NQSTY): Blue; Nonpolar (AILMFWV): Pink; Special (CGP): Black (B) Comparison of PeSA motifs created for different thresholds between 0.3 and 0.7, for known (p53K370, ErαK266, pRbK810, pRbK860 and HSB90K615) and predicted (SIX1K51, SIN3BK534 and DHX15K515) substrates by using the full motif, (C) and partial motif using the highest specificity positions. With a threshold value of 0.5, PeSA motif shows 80-100% match for all of the known substrates. 80% match corresponds to 2 out of 10 residues mismatch. Similar match success occurs for partial match as seen in (C).

Using the full motif comparison and allowing a single residue mismatch, PeSA motif would predict four of the five known substrates, failing to identify Rb (K860) from the array results. In contrast, the motif generated by Lanouette et al. also fails to identify Rb as a potential substrate at both K810 and K860 positions, as well as HSP90. As for the peptide sequences positively identified by Lanouette et al, PeSA motif successfully identifies SIX1 and DHX15 as potential substrates. By focusing PeSA analysis to the region of the motif displaying high specificity (i.e., residues −1 to +3), the similar trend can be seen for the predicted and known substrates tested. PeSA has 100% match for all known and predicted substrates, except Rb (K860) and SIN3B (K354). By allowing one mismatch as a threshold for substrate prediction, PeSA maintains a 100% success in re-predicting known and validated SMYD2 targets from peptide array results (Figure 5C).

## 4. CONCLUSION

Understanding protein-protein interactions, including substrate specificity of enzymes, has been a research question that has been attempted to be addressed by numerous studies (Fields and Song, 1989; Levy et al., 2011; Rathert et al., 2008; Wei et al., 2018). The utility of PeSA results from a design that allows the user to set experimental parameters and see the results instantly and effortlessly. Parameters include (1) the selection of different frequency or densitometry weight thresholds, or (2) number of residues per position. The ability to see how specific parameters alter the results will facilitate to make decisions on a per-case basis. PeSA also creates a base for a variety of future expansions. Motif is one form of visual output that can be used for data analysis. Filtering a list of possible peptide list for specificity by their similarity to a motif is the simplest form of utilizing a motif (Kudithipudi et al., 2014; Lanouette et al., 2015). Motifs are not merely end result of a study but can also be used as input data for other studies as well. Machine learning approach for prediction using validated motifs (Waardenberg et al., 2016) or finding relations between different motifs to create fingerprints (Attwood et al., 2000) are a couple of such examples. Similarly, the data generated by PeSA as the foundation of the motif can be used for many other purposes.

## ACKNOWLEDGEMENTS

This research was funded by an NSERC discovery grant (grant number 06151) to KK Biggar.

## REFERENCES

Arenkov, P., Kukhtin, A., Gemmell, A., Voloshchuk, S., Chupeeva, V., Mirzabekov, A., 2000. Protein microchips: Use for immunoassay and enzymatic reactions. Anal. Biochem. 278, 123–131. https://doi.org/10.1006/abio.1999.4363

Arifuzzaman, M., Maeda, M., Itoh, A., Nishikata, K., Takita, C., Saito, R., Ara, T., Nakahigashi, K., Huang, H., Hirai, A., Tsuzuki, K., Nakamura, S., Altaf-ul-amin, M., Oshima, T., Baba, T., Yamamoto, N., Kawamura, T., Ioka-nakamichi, T., Kitagawa, M., Tomita, M., Kanaya, S., Wada, C., Mori, H., 2006. Aristotle’s second thoughts.pdf. Genome Res. 16, 686–691. https://doi.org/10.1101/gr.4527806.8

Arrowsmith, C.H., Bountra, C., Fish, P. V., Lee, K., Schapira, M., 2012. Epigenetic protein families: a new frontier for drug discovery. Nat. Rev. 11, 384–400.

Attwood, T.K., Croning, M.D.R., Flower, D.R., Lewis, A.P., Mabey, J.E., Scordis, P., Selley, J.N., Wright, W., 2000. PRINTS-S□: the database formerly known as PRINTS 28, 225–227.

Baspinar, A., Cukuroglu, E., Nussinov, R., Keskin, O., Gursoy, A., 2014. PRISM: A web server and repository for prediction of protein-protein interactions and modeling their 3D complexes. Nucleic Acids Res. 42, 285–289. https://doi.org/10.1093/nar/gku397

Biggar, K.K., Li, S.S.C., 2014. Nonhistone_protein_methylation.pdf. Nat. Rev. Mol. Cell Biol. 16, 5–17.

Carpentier, G., 2010. Protein Array Analyzer. Available online: http://rsb.info.nih.gov/ij/macros/toolsets/ProteinArrayAnalyzer.txt

Creixell, P., Palmeri, A., Miller, C.J., Lou, H.J., Santini, C.C., Nielsen, M., Turk, B.E., Linding, R., 2015. Unmasking Determinants of Specificity in the Human Kinome. Cell 163, 187–201. https://doi.org/10.1016/j.cell.2015.08.057

Dhayalan, A., Kudithipudi, S., Rathert, P., Jeltsch, A., 2011. Specificity analysis-based identification of new methylation targets of the SET7/9 protein lysine methyltransferase. Chem. Biol. 18, 111–120. https://doi.org/10.1016/j.chembiol.2010.11.014

Fields, S., Song, O., 1989. A Novel Genetic System to Detect Protein-Protein Interactions. Nature 340, 245–246. https://doi.org/10.1038/340245a0

Franceschini, A., Szklarczyk, D., Frankild, S., Kuhn, M., Simonovic, M., Roth, A., Lin, J., Minguez, P., Bork, P., Von Mering, C., Jensen, L.J., 2013. STRING v9.1: Protein-protein interaction networks, with increased coverage and integration. Nucleic Acids Res. 41, 808–815. https://doi.org/10.1093/nar/gks1094

Ito, T., Chiba, T., Ozawa, R., Yoshida, M., Hattori, M., Sakaki, Y., 2001. A comprehensive two-hybrid analysis to explore the yeast protein interactome. Proc. Natl. Acad. Sci. U. S. A. 98, 4569–74. https://doi.org/10.1073/pnas.061034498

Källman, J., 2019. EPPlus. Available online: https://www.nuget.org/packages/EPPlus/

Kotlyar, M., Pastrello, C., Pivetta, F., Lo Sardo, A., Cumbaa, C., Li, H., Naranian, T., Niu, Y., Ding, Z., Vafaee, F., Broackes-Carter, F., Petschnigg, J., Mills, G.B., Jurisicova, A., Stagljar, I., Maestro, R., Jurisica, I., 2014. In silico prediction of physical protein interactions and characterization of interactome orphans. Nat. Methods 12, 79–84. https://doi.org/10.1038/nmeth.3178

Kudithipudi, S., Dhayalan, A., Kebede, A.F., Jeltsch, A., 2012. The SET8 H4K20 protein lysine methyltransferase has a long recognition sequence covering seven amino acid residues. Biochimie 94, 2212–2218. https://doi.org/10.1016/j.biochi.2012.04.024

Kudithipudi, S., Lungu, C., Rathert, P., Happel, N., Jeltsch, A., 2014. Substrate specificity analysis and novel substrates of the protein lysine methyltransferase NSD1. Chem. Biol. 21, 226–237. https://doi.org/10.1016/j.chembiol.2013.10.016

Lanouette, S., Davey, J.A., Elisma, F., Ning, Z., Figeys, D., Chica, R.A., Couture, J.F., 2015. Discovery of substrates for a SET domain lysine methyltransferase predicted by multistate computational protein design. Structure 23, 206–215. https://doi.org/10.1016/j.str.2014.11.004

Lassle, M., Blatch, G.L., 1999. The tetratricopeptide repeat: a structural motif mediating protein-protein interactions. Bioessays 21, 932–9.

Levy, D., Liu, C.L., Yang, Z., Newman, A.M., Alizadeh, A.A., Utz, P.J., Gozani, O., 2011. A proteomic approach for the identification of novel lysine methyltransferase substrates. Epigenetics Chromatin 4, 1–12.

Li, N., Yu, Z., Ji, Q., Sun, J., Liu, X., Du, M., Zhang, W., 2017. An enzyme-mediated protein-fragment complementation assay for substrate screening of sortase A. Biochem. Biophys. Res. Commun. 486, 257–263. https://doi.org/10.1016/j.bbrc.2017.03.016

Li, P., Wang, L., Di, L., 2019. Applications of Protein Fragment Complementation Assays for Analyzing Biomolecular Interactions and Biochemical Networks in Living Cells. J. Proteome Res. acs.jproteome.9b00154. https://doi.org/10.1021/acs.jproteome.9b00154

Liu, H., Galka, M., Mori, E., Liu, X., Lin, Y. fen, Wei, R., Pittock, P., Voss, C., Dhami, G., Li, X., Miyaji, M., Lajoie, G., Chen, B., Li, S.S.C., 2013. A method for systematic mapping of protein lysine methylation identifies functions for HP1β in DNA damage response. Mol. Cell 50, 723–735. https://doi.org/10.1016/j.molcel.2013.04.025

Murakami, Y., Tripathi, L.P., Prathipati, P., Mizuguchi, K., 2017. Network analysis and in silico prediction of protein–protein interactions with applications in drug discovery. Curr. Opin. Struct. Biol. 44, 134–142. https://doi.org/10.1016/j.sbi.2017.02.005

Newton-King, J., 2019. Newtonsoft.Json. Available online: https://github.com/JamesNK

Pitre, S., Dehne, F., Chan, A., Cheetham, J., Duong, A., Emili, A., Gebbia, M., Greenblatt, J., Jessulat, M., Krogan, N., Luo, X., Golshani, A., 2006. PIPE: A protein-protein interaction prediction engine based on the re-occurring short polypeptide sequences between known interacting protein pairs. BMC Bioinformatics 7, 1–15. https://doi.org/10.1186/1471-2105-7-365

Primeau, M., Trudeau, L., Michnick, S.W., 2002. β-Lactamase protein fragment complementation assays as. Nat. Biotechnol. 20, 619–622.

Rasband, W., 2015. ImageJ. Available online: https://imagej.nih.gov/ij/

Rathert, P., Zhang, X., Freund, C., Cheng, X., Jeltsch, A., 2008. Analysis of the Substrate Specificity of the Dim-5 Histone Lysine Methyltransferase Using Peptide Arrays. Chem. Biol. 15, 5–11. https://doi.org/10.1016/j.chembiol.2007.11.013

Rayasam, G.V., Wendling, O., Angrand, P.O., Mark, M., Niederreither, K., Song, L., Lerouge, T., Hager, G.L., Chambon, P., Losson, R., 2003. NSD1 is essential for early post-implantation development and has a catalytically active SET domain. EMBO J. 22, 3153–3163. https://doi.org/10.1093/emboj/cdg288

Rolland, T., Taşan, M., Charloteaux, B., Pevzner, S.J., Zhong, Q., Sahni, N., Yi, S., Lemmens, I., Fontanillo, C., Mosca, R., Kamburov, A., Ghiassian, S.D., Yang, X., Ghamsari, L., Balcha, D., Begg, B.E., Braun, P., Brehme, M., Broly, M.P., Carvunis, A.-R., Convery-Zupan, D., Corominas, R., Coulombe-Huntington, J., Dann, E., Dreze, M., Dricot, A., Fan, C., Franzosa, E., Gebreab, F., Gutierrez, B.J., Hardy, M.F., Jin, M., Kang, S., Kiros, R., Lin, G.N., Luck, K., MacWilliams, A., Menche, J., Murray, R.R., Palagi, A., Poulin, M.M., Rambout, X., Rasla, J., Reichert, P., Romero, V., Ruyssinck, E., Sahalie, J.M., Scholz, A., Shah, A.A., Sharma, A., Shen, Y., Spirohn, K., Tam, S., Tejeda, A.O., Trigg, S.A., Twizere, J.-C., Vega, K., Walsh, J., Cusick, M.E., Xia, Y., Barabási, A.-L., Iakoucheva, L.M., Aloy, P., De Las Rivas, J., Tavernier, J., Calderwood, M.A., Hill, D.E., Hao, T., Roth, F.P., Vidal, M., 2014. A proteome-scale map of the human interactome network. Cell 159, 1212–1226. https://doi.org/10.1016/j.cell.2014.10.050

Rual, J.-F., Venkatesan, K., Hao, T., Hirozane-Kishikawa, T., Dricot, A., Li, N., Berriz, G.F., Gibbons, F.D., Dreze, M., Ayivi-Guedehoussou, N., Klitgord, N., Simon, C., Boxem, M., Milstein, S., Rosenberg, J., Goldberg, D.S., Zhang, L. V, Wong, S.L., Franklin, G., Li, S., Albala, J.S., Lim, J., Fraughton, C., Llamosas, E., Cevik, S., Bex, C., Lamesch, P., Sikorski, R.S., Vandenhaute, J., Zoghbi, H.Y., Smolyar, A., Bosak, S., Sequerra, R., Doucette-Stamm, L., Cusick, M.E., Hill, D.E., Roth, F.P., Vidal, M., 2005. Towards a proteome-scale map of the human protein-protein interaction network. Nature 437, 1173–8. https://doi.org/10.1038/nature04209

Schuhmacher, M.K., Kudithipudi, S., Kusevic, D., Weirich, S., Jeltsch, A., 2015. Activity and specificity of the human SUV39H2 protein lysine methyltransferase. Biochim. Biophys. Acta - Gene Regul. Mech. 1849, 55–63. https://doi.org/10.1016/j.bbagrm.2014.11.005

Uetz, P., Giot, L., Cagney, G., Mansfield, T.A., Judson, R.S., Knight, J.R., Lockshon, D., Narayan, V., Srinivasan, M., Pochart, P., Qureshi-Emili, A., Li, Y., Godwin, B., Conover, D., Kalbfleisch, T., Vijayadamodar, G., Yang, M., Johnston, M., Fields, S., Rothberg, J.M., 2000. A comprehensive analysis of protein-protein interactions in Saccharomyces cerevisiae. Nature 403, 623–627. https://doi.org/10.1038/35001009

Waardenberg, A.J., Homan, B., Mohamed, S., Harvey, R.P., Bouveret, R., Harvey, R.P., 2016. Prediction and validation of protein – protein interactors from genome-wide DNA-binding data using a knowledge-based machine-learning approach. Open Biol. 6, 160183.

Wei, R., Kaneko, T., Liu, X., Liu, H., Li, L., Voss, C., Liu, E., He, N., Li, S.S.-C., 2018. Interactome Mapping Uncovers a General Role for Numb in Protein Kinase Regulation. Mol. Cell. Proteomics 17, 2216–2228. https://doi.org/10.1074/mcp.ra117.000114

Weimann, M., Grossmann, A., Woodsmith, J., Özkan, Z., Birth, P., Meierhofer, D., Benlasfer, N., Valovka, T., Timmermann, B., Wanker, E.E., Sauer, S., Stelzl, U., 2013. A Y2H-seq approach defines the human protein methyltransferase interactome. Nat. Methods 10, 339–342.

Weirich, S., Kudithipudi, S., Jeltsch, A., 2016. Specificity of the SUV4-20H1 and SUV4-20H2 protein lysine methyltransferases and methylation of novel substrates. J. Mol. Biol. 428, 2344–2358. https://doi.org/10.1016/j.jmb.2016.04.015

Yu, X., Noll, R.R., Romero Dueñas, B.P., Allgood, S.C., Barker, K., Caplan, J.L., Machner, M.P., LaBaer, J., Qiu, J., Neunuebel, M.R., 2018. Legionella effector AnkX interacts with host nuclear protein PLEKHN1. BMC Microbiol. 18, 1–14. https://doi.org/10.1186/s12866-017-1147-7

Zhu, G., Liu, Y., Shaw, S., 2005. Protein Kinase Specificity: A Strategic Collaboration between Kinase Peptide Specificity and Substrate Recruitment. Cell Cycle 4, 52–56.

